# Non-invasive stimulation of vagal afferents reduces gastric frequency

**DOI:** 10.1101/773259

**Authors:** Vanessa Teckentrup, Sandra Neubert, João C. P. Santiago, Manfred Hallschmid, Martin Walter, Nils B. Kroemer

**Affiliations:** University of Tübingen, Department of Psychiatry and Psychotherapy, Germany; University of Tübingen, Department of Medical Psychology and Behavioral Neurobiology, Germany; German Center for Diabetes Research (DZD), Tübingen, Germany; Institute for Diabetes Research and Metabolic Diseases of the Helmholtz Center Munich at the Eberhard Karls University Tübingen, Tübingen, Germany; University of Magdeburg, Department of Psychiatry and Psychotherapy, Germany; Leibniz Institute for Neurobiology, Magdeburg, Germany; University of Jena, Department of Psychiatry and Psychotherapy, Germany

**Author notes:** **Corresponding authors**, Vanessa Teckentrup, Dr. Nils B. Kroemer, Calwerstr. 14, 72076 Tübingen, Germany.

## Abstract

Metabolic feedback between the gut and the brain relayed via the vagus nerve contributes to energy homeostasis. We investigated in healthy adults whether non-invasive stimulation of vagal afferents impacts energy homeostasis via efferent effects on metabolism or digestion. In a randomized crossover design, we applied transcutaneous auricular vagus nerve stimulation (taVNS) while recording efferent metabolic effects using simultaneous electrogastrography (EGG) and indirect calorimetry. We found that taVNS reduced gastric myoelectric frequency (*p* =.008), but did not alter resting energy expenditure. We conclude that stimulating vagal afferents induces gastric slowing via vagal efferents without acutely affecting net energy expenditure at rest. Collectively, this highlights the potential of taVNS to modulate digestion by activating the dorsal vagal complex. Thus, taVNS-induced changes in gastric frequency are an important peripheral marker of brain stimulation effects.

## 1. Introduction

Maintaining energy homeostasis is vital for organisms and necessitates a balance between energy intake and expenditure [1]. Achieving this balance requires vagal afferents to transmit information between peripheral organs and the dorsal vagal complex in the brain stem [2–6]. Invasive stimulation of the vagus nerve (VNS) as well as the more recent non-invasive transcutaneous auricular VNS (taVNS, [7–9]) impact energy homeostasis by modulating food intake, energy metabolism, and glycemic control [10–12]. In rodents, VNS triggered by phasic stomach contractions resulted in weight loss [13]. In humans, taVNS led to a decreased frequency and increased amplitude of gastric motility [14]. Such metabolic effects might be related to VNS-induced increases in the activity of brown adipose tissue, which in turn increased the basal metabolic rate [15]. Notably, dopamine has been suggested as a neuromodulator of energy homeostasis within the gut-brain axis [16,17]. Afferently, stimulation of the vagal sensory ganglion in mice was found to induce dopamine release in the substantia nigra [17]. Efferently, dopamine administration to the dorsal vagal complex in rats modulated the upper gastrointestinal tract by reducing gastric tone and motility via DA2 receptors in the dorsal motor nucleus of the vagus [18]. Thus, while vagal stimulation mostly targets afferent pathways, studies in rodents provide evidence for brain-mediated effects on downstream targets.

Although there is preliminary evidence linking vagal signaling and energy homeostasis [14], efferent taVNS-induced effects on digestion and energy metabolism in healthy humans have not been conclusively demonstrated. We therefore investigated whether taVNS vs. sham changes electrogastrography (EGG) and indirect calorimetry as markers of energy homeostasis.

## 2. Methods

### 2.1 Participants and procedure

We included 22 participants (14 female, M_age_ ± SD = 23.3 ± 2.7 years, range: 19-29) in the study. In a randomized crossover design, we measured EGG using four standard electrocardiogram electrodes connected to a BrainProducts BrainAmp DC EEG recording system. Electrodes were placed as previously described [19]. Resting energy expenditure (REE) was measured with the CareFusion Vmax ventilated hood system for indirect calorimetry (see SI). For administering taVNS, we used Cerbomed NEMOS following the protocol presented in ref. [8]. Briefly, the electrode was placed at the left cymba conchae (taVNS) or was turned upside down and placed at the earlobe (sham). The stimulation protocol of NEMOS is preset with a biphasic impulse frequency of 25 Hz with alternating intervals of 30 s stimulation on and 30 s off.

After a resting period of at least 15 minutes, we recorded a 15-minute baseline for both EGG and calorimetry. Next, we placed the taVNS device on the participants’ left ear according to the randomization protocol. The individual stimulation intensity was adjusted based on subjective pain thresholds using concurrent VAS ratings (for details, see [20]). We then recorded at least 30 minutes of EGG and calorimetry during active stimulation before the participant was debriefed.

### 2.2 Data preprocessing and statistical analysis

EGG data were preprocessed and inspected for muscle artifacts. We then identified the gastric peak frequency for baseline, taVNS and sham, respectively, based on spectral density for each EGG channel (see SI). One participant had to be excluded after quality control due to absence of visibly identifiable peaks in any channel in both sessions, leaving N=21 for the statistical analysis.

We calculated baseline-corrected delta mean gastric frequency (in mHz) by subtracting the individual session-specific baseline mean gastric frequency from the respective taVNS and sham mean gastric frequency. Next, we calculated the net effect of stimulation (interaction) by subtracting delta sham from delta taVNS. After preprocessing the calorimetry data (see SI), we calculated the same measures for REE (in kcal/day). For non-parametric inference, we bootstrapped the distribution of taVNS-induced changes in gastric frequency and REE, respectively, using 50,000 repetitions and calculated two-tailed p-values.

## 3. Results

We found that taVNS compared to sham led to a significant reduction in gastric myoelectric frequency (*Figure 1A;* mean [95% bootstrap CI] Time × Stimulation: −2.24 mHz [−4.44, −0.72], *p_boot_*= .008). In contrast, we observed no significant effect of taVNS on resting energy expenditure (*Figure 1B;* mean [95% bootstrap CI] Time × Stimulation: −3.69 kcal/day [−46.52, 42.31], *p_boot_*= .863).

**Figure 1.**
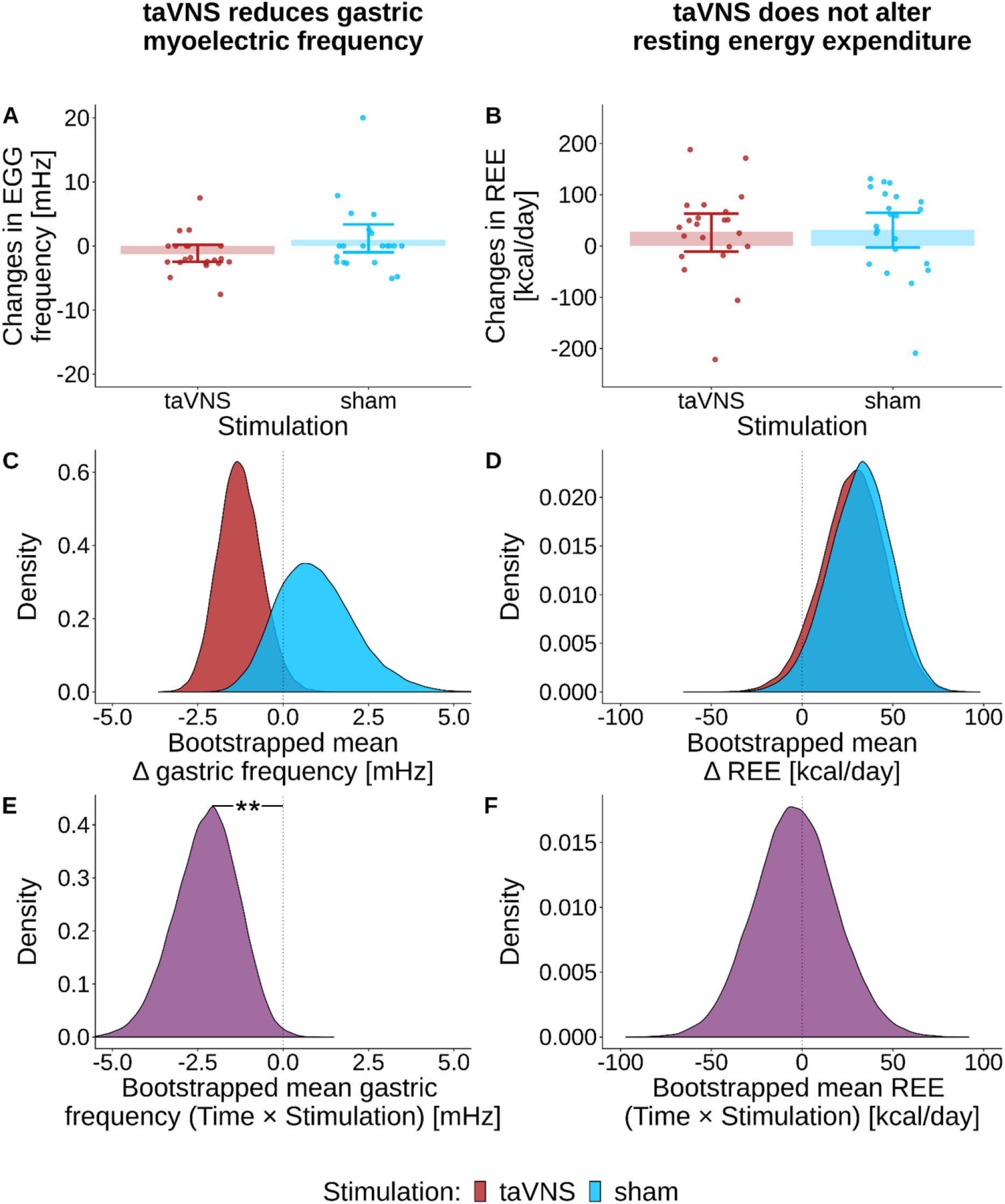
Gastric frequency is reduced after transcutaneous auricular vagus nerve stimulation (taVNS) compared to sham stimulation, but there is no change in resting energy expenditure (REE). The figure depicts A) changes (stimulation – baseline) in gastric frequency and B) changes (stimulation – baseline) in REE for taVNS and sham. It further depicts C) bootstrapped delta gastric frequency distributions (stimulation – baseline) and D) bootstrapped delta REE distributions (stimulation – baseline) as well as the interaction between Time (pre, post) and Stimulation (taVNS, sham) for E) EGG and F) REE. While the interaction is significant for gastric frequency with pboot = .008, the same term does not reach significance for REE with pboot = 863. Bootstrapped distributions are based on 50,000 iterations.

## 4. Discussion

In line with the hypothesized efferent effect, we found that taVNS alters a marker of energy homeostasis in humans. The observed taVNS-induced reduction in gastric frequency is well in line with previous findings linking VNS to altered energy homeostasis [13,14,17]. This efferent effect on gastric motility might be due to a taVNS-induced release of dopamine in the brain stem. Previous work has shown that elevated levels of brain stem dopamine lead to reduced food intake [21] and gastric relaxation [22]. Moreover, dopamine administration in the brain stem reduced gastric tone and motility which was abolished by vagotomy [18]. Studies linking alterations in vagal signaling to the development of Parkinson’s disease [23,24] further support the assumption of afferent signaling between the gut and key dopaminergic brain regions along the vagal pathway. Therefore, taVNS-induced neuromodulation in the brain stem might lead to the observed slowing of gastric myoelectric frequency via the efferent vagal pathway.

In contrast to chronic VNS in patients [15], we did not find changes in energy expenditure during acute taVNS. This pattern indicates that compared to changes in digestion taVNS-induced effects on energy expenditure may develop across longer time periods.

In sum, we demonstrated that taVNS reduces gastric frequency without affecting REE. This shows that transcutaneous stimulation of vagal afferents can elicit efferent gastric effects through a feedback loop via the dorsal vagal complex. Thus, in light of the heterogeneous efferent effects of taVNS on electrocardiogram parameters [25,26], the EGG may be a promising positive control measure for taVNS in healthy humans [27].

## Acknowledgements

We thank Caroline Burrasch, Franziska Müller, Moritz Herkner and Julius Gervelmeyer for help with data acquisition. This study was supported by the University of Tübingen, Faculty of Medicine fortüne grant #2453-0-0. NBK received support from the Else Kröner-Fresenius-Stiftung, grant #2017-A67. JCPS and MH received support via grants from the German Federal Ministry of Education and Research (BMBF) to the German Center for Diabetes Research (DZD e.V.; 01GI0925).

## Author contributions

NBK was responsible for the concept and design of the study. VT & JCPS collected data. NBK & VT conceived the method and processed the data. VT performed the data analysis and NBK & SN contributed to analyses. VT, SN & NBK wrote the manuscript. All authors contributed to the interpretation of findings, provided critical revision of the manuscript for important intellectual content and approved the final version for publication.

## References

[1] Waterson MJ, Horvath TL. Neuronal Regulation of Energy Homeostasis: Beyond the Hypothalamus and Feeding. Cell Metab 2015;22:962–70. doi:10.1016/j.cmet.2015.09.026.

[2] Gribble FM, Reimann F. Function and mechanisms of enteroendocrine cells and gut hormones in metabolism. Nat Rev Endocrinol 2019;15:226–37. doi:10.1038/s41574-019-0168-8.

[3] Small DM, DiFeliceantonio AG. Processed foods and food reward. Science 2019;363:346–7. doi:10.1126/science.aav0556.

[4] de Lartigue G. Role of the vagus nerve in the development and treatment of diet-induced obesity. J Physiol 2016;594:5791–815. doi:10.1113/JP271538.

[5] Johnson RL, Wilson CG. A review of vagus nerve stimulation as a therapeutic intervention. J Inflamm Res 2018;11:203–13. doi:10.2147/JIR.S163248.

[6] Mizrahi M, Ben Ya’acov A, Ilan Y. Gastric stimulation for weight loss. World J Gastroenterol 2012;18:2309–19. doi:10.3748/wjg.v18.i19.2309.

[7] Fallgatter AJ, Neuhauser B, Herrmann MJ, Ehlis A-C, Wagener A, Scheuerpflug P, et al. Far field potentials from the brain stem after transcutaneous vagus nerve stimulation. J Neural Transm 2003;110:1437–43. doi:10.1007/s00702-003-0087-6.

[8] Frangos E, Ellrich J, Komisaruk BR. Non-invasive Access to the Vagus Nerve Central Projections via Electrical Stimulation of the External Ear: fMRI Evidence in Humans. Brain Stimul 2015;8:624–36. doi:10.1016/j.brs.2014.11.018.

[9] Yakunina N, Kim SS, Nam E-C. Optimization of Transcutaneous Vagus Nerve Stimulation Using Functional MRI. Neuromodulation 2017;20:290–300. doi:10.1111/ner.12541.

[10] Cork SC. The role of the vagus nerve in appetite control: Implications for the pathogenesis of obesity. J Neuroendocrinol 2018;30:e12643. doi:10.1111/jne.12643.

[11] Ikramuddin S, Blackstone RP, Brancatisano A, Toouli J, Shah SN, Wolfe BM, et al. Effect of reversible intermittent intra-abdominal vagal nerve blockade on morbid obesity: the ReCharge randomized clinical trial. JAMA 2014;312:915–22. doi:10.1001/jama.2014.10540.

[12] Shikora S, Toouli J, Herrera MF, Kulseng B, Zulewski H, Brancatisano R, et al. Vagal blocking improves glycemic control and elevated blood pressure in obese subjects with type 2 diabetes mellitus. J Obes 2013;2013:245683. doi:10.1155/2013/245683.

[13] Yao G, Kang L, Li J, Long Y, Wei H, Ferreira CA, et al. Effective weight control via an implanted self-powered vagus nerve stimulation device. Nat Commun 2018;9:5349. doi:10.1038/s41467-018-07764-z.

[14] Hong G-S, Pintea B, Lingohr P, Coch C, Randau T, Schaefer N, et al. Effect of transcutaneous vagus nerve stimulation on muscle activity in the gastrointestinal tract (transVaGa): a prospective clinical trial. Int J Colorectal Dis 2018. doi:10.1007/s00384-018-3204-6.

[15] Vijgen GHEJ, Bouvy ND, Leenen L, Rijkers K, Cornips E, Majoie M, et al. Vagus nerve stimulation increases energy expenditure: relation to brown adipose tissue activity. PLoS One 2013;8:e77221. doi:10.1371/journal.pone.0077221.

[16] Tellez LA, Medina S, Han W, Ferreira JG, Licona-Limón P, Ren X, et al. A gut lipid messenger links excess dietary fat to dopamine deficiency. Science 2013;341:800–2. doi: 10.1126/science.1239275.

[17] Han W, Tellez LA, Perkins MH, Perez IO, Qu T, Ferreira J, et al. A Neural Circuit for Gut-Induced Reward. Cell 2018;175:887–8. doi:10.1016/j.cell.2018.10.018.

[18] Anselmi L, Toti L, Bove C, Travagli RA. Vagally mediated effects of brain stem dopamine on gastric tone and phasic contractions of the rat. Am J Physiol Gastrointest Liver Physiol 2017;313:G434–41. doi:10.1152/ajpgi.00180.2017.

[19] Rebollo I, Devauchelle A-D, Béranger B, Tallon-Baudry C. Stomach-brain synchrony reveals a novel, delayed-connectivity resting-state network in humans. Elife 2018;7. doi:10.7554/eLife.33321.

[20] Kuehnel A, Teckentrup V, Neuser MP, Huys QJM, Burrasch C, Walter M, et al. Stimulation of the vagus nerve reduces learning in a go/no-go reinforcement learning task. bioRxiv 2019. doi:10.1101/535260.

[21] Kaplan JM, Södersten P. Apomorphine suppresses ingestive behaviour in chronic decerebrate rats. Neuroreport 1994;5:1839–40. doi:10.1097/00001756-199409080-00039.

[22] Valenzuela JE. Dopamine as A Possible Neurotransmitter in Gastric Relaxation. Gastroenterology 1976;71:1019–22. doi:10.1016/S0016-5085(76)80051-0.

[23] Svensson E, Horváth-Puhó E, Thomsen RW, Djurhuus JC, Pedersen L, Borghammer P, et al. Vagotomy and subsequent risk of Parkinson’s disease. Ann Neurol 2015;78:522–9. doi:10.1002/ana.24448.

[24] Liu B, Fang F, Pedersen NL, Tillander A, Ludvigsson JF, Ekbom A, et al. Vagotomy and Parkinson disease: A Swedish register-based matched-cohort study. Neurology 2017;88:1996–2002. doi:10.1212/WNL.0000000000003961.

[25] De Couck M, Cserjesi R, Caers R, Zijlstra WP, Widjaja D, Wolf N, et al. Effects of short and prolonged transcutaneous vagus nerve stimulation on heart rate variability in healthy subjects. Auton Neurosci 2017;203:88–96. doi:10.1016/j.autneu.2016.11.003.

[26] Bauer S, Baier H, Baumgartner C, Bohlmann K, Fauser S, Graf W, et al. Transcutaneous Vagus Nerve Stimulation (tVNS) for Treatment of Drug-Resistant Epilepsy: A Randomized, Double-Blind Clinical Trial (cMPsE02). Brain Stimul 2016;9:356–63. doi:10.1016/j.brs.2015.11.003.

[27] Teckentrup V, Krylova M, Jamalabadi H, Neubert S, Neuser MP, Hartig R, et al. Brain signaling dynamics after vagus nerve stimulation. NeuroImage 2019. In-principle accepted stage 1 registered report.

