## Supplementary Information for "Non-invasive stimulation of vagal afferents reduces gastric frequency"

Vanessa Teckentrup[1*], Sandra Neubert[1], João C. P. Santiago[2,3,4], Manfred Hallschmid[2,3,4], Martin Walter[1,5,6,7], Nils B. Kroemer[1*]

1 University of Tübingen, Department of Psychiatry and Psychotherapy, Germany
2 University of Tübingen, Department of Medical Psychology and Behavioral Neurobiology, Germany
3 German Center for Diabetes Research (DZD), Tübingen, Germany
4 Institute for Diabetes Research and Metabolic Diseases of the Helmholtz Center Munich at the Eberhard Karls University Tübingen, Tübingen, Germany
5 University of Magdeburg, Department of Psychiatry and Psychotherapy, Germany
6 Leibniz Institute for Neurobiology, Magdeburg, Germany
7 University of Jena, Department of Psychiatry and Psychotherapy, Germany

**Corresponding authors***

Vanessa Teckentrup,

Dr. Nils B. Kroemer,

Calwerstr. 14, 72076 Tübingen, Germany

### 1. Methods

#### 1.1 Participants and procedure

Both experimental days were scheduled back to back and participants arrived between 7:00 and 12:30 am. They were instructed not to consume any calories for at least 4 hours and not to drink anything for 2 hours prior to the experiment. Moreover, they were not allowed to engage in strenuous physical activity 24 hours prior to the experiment. After providing written informed consent, participants were asked regarding their last meal and drink as well as their physical activity on the day of the measurement. We furthermore measured weight, height and waist and hip circumference. Participants were then asked to lie down on a bed. To measure the electrogastrogram (EGG), the skin above the abdomen was cleaned and six (four EGG electrodes, reference and ground electrode) standard electrocardiogram electrodes with solid gel (3M red dot) were applied according to the placement described in [1]. The electrodes were connected to a BrainAmp DC (BrainProducts, Gilching, Germany) EEG recording system and electrodes were left in place for the second session. EGG was acquired at a sampling rate of 250 Hz with a low-pass filter of 80 Hz.

To assess resting energy expenditure (REE), we used the Vmax (CareFusion, San Diego, CA, USA) ventilated hood system for indirect calorimetry which measures the ratios of $CO_2$ and $O_2$ flowing in vs. out of the ventilated hood. During measurement, participants were asked to lie still on the bed and breathe normally.

For administering taVNS, we used Cerbomed NEMOS (Erlangen, Germany) following the protocol of [2]. The electrode was placed at the left cymba conchae (taVNS) or was turned upside down and placed at the earlobe (sham). Prior to each measurement, the stimulus intensity was individually adjusted based on subjective pain thresholds using concurrent VAS ratings [3]. As recommended by the manufacturer, the intensity was increased from 0.1 mA in 0.1 mA increments until participants reported a “tingling” sensation (which was supposed to not be painful). Given this matching procedure, participants do not guess better than chance which stimulation condition they had received ([4], total of recorded guesses: 148, correct guesses: 79, accuracy: 53.4%, $p_{binom}$ = .18). The stimulation protocol of NEMOS is preset with a biphasic impulse frequency of 25 Hz with alternating intervals of 30 seconds stimulation on and 30 seconds off.

During the entire recording, we played an audiobook for the participants in order to keep them from falling asleep. After a minimum resting period of 15 minutes during which the indirect calorimetry device was calibrated, we started with a 15-minute baseline measurement for both EGG and calorimetry. Next, we placed the taVNS device on the participants' left ear according to the randomization protocol and adjusted the stimulus intensity as described above. We then recorded at least 30 minutes of EGG and indirect calorimetry with active stimulation before the participant was debriefed. Markers for EGG were set manually and coded for the removal and respective re-application of the calorimetry hood during placement of the NEMOS earpiece, the start of the stimulation phase and any disturbances during the recording (such as loss of contact of the NEMOS earpiece, calorimetry device malfunction or excessive movement of the subject).

#### 1.2 Data preprocessing and statistical analysis

Data from EGG was read into MATLAB R2017a (The Mathworks Inc., Natick, MA, USA) and processed using fieldtrip [5] based on scripts released by [1][1]. We cut the data according to the markers set during measurement to get two EGG time series, one for the baseline phase and one for the stimulation phase (taVNS or sham). Next, we low-pass filtered the data at 5 Hz to prevent aliasing, downsampled to 10 Hz and the mean was removed from the time series. For quality control, the time series were visually inspected for muscle artifacts. Affected segments were marked and only clean, continuous segments were kept for further analyses. To identify the gastric peak frequency for baseline, taVNS and sham, respectively, we used the fast fourier transform with a Hanning taper to calculate spectral density for each EGG channel. The peak and associated EGG channel were then identified via visual inspection

[1] https://github.com/irebollo/stomach_brain_Scripts

based on sharpness and power within the frequency range of interest (0.033 - 0.066 Hz, *Table S1*). For calorimetry data, we excluded the first 3 minutes after placing the hood to allow the system to reach equilibrium and extracted the measure of mean REE (in kcal/day) for statistical analysis (*Table S1*). For one participant, the REE baseline measurement for the first session (sham condition) was not available due to a technical error. This was compensated for by using the respective participants' REE baseline measurement from the second session.

**Table S1.** Mean gastric peak frequency (in Hz) as well as mean resting energy expenditure (in kcal/day) and associated standard deviations for the two baseline measurements, taVNS and sham.

| | Baseline taVNS | Baseline sham | Stimulation taVNS | Stimulation sham |
|---|---|---|---|---|
| Gastric frequency [Hz ± SD] | 0.0484 ± 0.004 | 0.0488 ± 0.006 | 0.0471 ± 0.003 | 0.0498 ± 0.005 |
| Resting energy expenditure [kcal/day ± SD] | 1491.187 ± 367.087 | 1489.06 ± 339.95 | 1519.009 ± 312.488 | 1520.555 ± 333.533 |

### References

[1] Rebollo I, Devauchelle A-D, Béranger B, Tallon-Baudry C. Stomach-brain synchrony reveals a novel, delayed-connectivity resting-state network in humans. Elife 2018;7. doi:10.7554/eLife.33321.

[2] Frangos E, Ellrich J, Komisaruk BR. Non-invasive Access to the Vagus Nerve Central Projections via Electrical Stimulation of the External Ear: fMRI Evidence in Humans. Brain Stimul 2015;8:624–36. doi:10.1016/j.brs.2014.11.018.

[3] Kuehnel A, Teckentrup V, Neuser MP, Huys QJM, Burrasch C, Walter M, et al. Stimulation of the vagus nerve reduces learning in a go/no-go reinforcement learning task. bioRxiv 2019. doi:10.1101/535260.

[4] Neuser MP, Teckentrup V, Kuehnel A, Hallschmid M, Walter M, Kroemer NB. Vagus nerve stimulation increases vigor to work for rewards. Unpublished results.

[5] Oostenveld R, Fries P, Maris E, Schoffelen J-M. FieldTrip: Open source software for advanced analysis of MEG, EEG, and invasive electrophysiological data. Comput Intell Neurosci 2011;2011:156869. doi:10.1155/2011/156869.
